# Leopard: fast decoding cell type-specific transcription factor binding landscape at single-nucleotide resolution

**DOI:** 10.1101/856823

**Authors:** Hongyang Li, Yuanfang Guan

**Affiliations:** Department of Computational Medicine and Bioinformatics, University of Michigan, 100 Washtenaw Avenue, Ann Arbor, MI 48109, USA

**Author notes:** Email addresses: Hongyang Li, Yuanfang Guan.

**Keywords:** transcription factor binding site, single-nucleotide resolution, neural network, deep learning, ChIP-seq, ChIP-exo, DNase-seq

## Abstract

Decoding the cell type-specific transcription factor (TF) binding landscape at single-nucleotide resolution is crucial for understanding the regulatory mechanisms underlying many fundamental biological processes and human diseases. However, limits on time and resources restrict the high-resolution experimental measurements of TF binding profiles of all possible TF-cell type combinations. Previous computational approaches either can not distinguish the cell-context-dependent TF binding profiles across diverse cell types, or only provide a relatively low-resolution prediction. Here we present a novel deep learning approach, Leopard, for predicting TF-binding sites at single-nucleotide resolution, achieving the median area under receiver operating characteristic curve (AUROC) of 0.994. Our method substantially outperformed state-of-the-art methods Anchor and FactorNet, improving the performance by 19% and 27% respectively despite evaluated at a lower resolution. Meanwhile, by leveraging a many-to-many neural network architecture, Leopard features hundred-fold to thousand-fold speedup compared to current many-to-one machine learning methods.

## Background

Transcription factors (TFs) play a fundamental role in regulating gene expression via binding to specific DNA sequences [1–3]. Precisely decoding the transcription factor binding landscape at single-nucleotide resolution is crucial for understanding the regulatory mechanisms underlying many cellular processes and human diseases [4–7]. Beyond the sequence preferences of TF binding represented as motifs [8,9], the Encyclopedia of DNA elements (ENCODE) project has established that TFs almost exclusively bind to open chromatin [10,11]. The TF binding landscapes therefore vary substantially across cell types, which are associated with their unique organization of accessible chromatin across the genome [10,12]. Chromatin immunoprecipitation followed by DNA sequencing (ChIP-seq) is a common technique to measure the *in vivo* TF binding profile in a specific cell type [13]. Unfortunately, the contamination of immunoprecipitations from unbound DNA in ChIP-seq experiments leads to the description of TF binding only at a relatively low resolution [5,14]. An alternative approach, ChIP-exo, can precisely map the genome-wide TF binding locations and reduce the erroneous calls [5,15]. However, it is infeasible to experimentally measure the single-nucleotide-resolution TF binding landscapes in the enormous combinations of TF and cell type pairs due to limits on time and resources.

Recent advancements in computational models show great promise to delineate the TF binding landscape *in silico [16,17]*. Previous pioneering works including DeepSea [18], DeepBind [19], DanQ [20], and DeFine [21] have modeled the relationships between DNA sequences and TFs using neural network models. However, without the cell type-specific information on chromatin accessibility, these models can not distinguish the diverse TF binding profiles across different cell types and conditions. Recent methods including Anchor [22] and FactorNet [23] address this problem by considering both DNA sequence and chromatin accessibility, greatly improving the prediction accuracy in a cross-cell type fashion. But these methods typically only provide the statistically enriched TF binding regions at around 200 base pair (bp) resolution. Therefore, there is a great demand for computational tools to both accurately and precisely model the TF binding status for every single genomic position.

Here we present a novel deep neural network approach, Leopard, for decoding cell type-specific TF binding landscapes at single-nucleotide resolution. Using both DNA sequence and chromatin accessibility from DNase-seq as inputs, Leopard accurately predict genome-wide TF binding locations with the median area under receiver operating characteristic curve (AUROC) of 0.994 on a large held-out testing dataset of 27 ChIP-seq profiles. When benchmarked against ChIP-exo experiment results, our method identified single-nucleotide TF binding peaks although it was only trained on low-resolution ChIP-seq data. Compared with state-of-the-art methods that make prediction at 200bp resolution, Leopard substantially surpassed Anchor and FactorNet, improving the performance by 19% and 27% respectively in 12 TFs under consideration. Similar to computer vision models for pixel-level image segmentation, a many-to-many nucleotide-level segmentation framework was leveraged to generate predictions for multiple genomic positions simultaneously, allowing for hundred-fold to thousand-fold speedup compared with the current many-to-one deep learning and traditional machine learning models. In summary, Leopard features precise, accurate and fast single-base decoding of TF binding landscapes across different cell types.

## Results

### Overview of experimental design

Leopard is designed to predict cross-cell type TF binding sites at single-nucleotide level based on DNA sequence and chromatin accessibility from DNase-seq (Fig. 1). A total of 69 ChIP-seq experiments and 13 DNase-seq experiments were used from the ENCODE project, covering 12 TFs (CTCF, E2F1, EGR1, FOXA1, FOXA2, GABPA, HNF4A, JUND, MAX, NANOG, REST, and TAF1) in 13 cell types (A549, GM12878, H1-hESC, HCT116, HeLa-S3, HepG2, iPSC, IMR-90, K562, liver, MCF-7, Panc1, PC-3). For each TF-cell type pair, we used the real-valued DNase-seq filtered alignment signal as the primary feature. To alleviate the potential sequencing biases and batch effects, we calculated the ΔDNase-seq signal by subtracting the average signal across all 13 cell types under consideration [22]. To capture TF binding motifs, DNA sequences were one-hot encoded and used as additional input features. For the training labels, instead of directly using ChIP-seq peaks, we refined these relatively low-resolution peaks by overlapping them with FIMO motif scanning signals [24]. A deep many-to-many convolutional neural network model was designed to model the cell type-specific contexts and DNA sequences, identifying TF binding sites at single-nucleotide resolution. We used a nested train-validate-test framework, where the parameters of the neural network were learned on the training data and the hyper-parameters were tuned on the validation data. A subset of 27 ChIP-seq data were held out as the testing set to evaluate the performance of our model (Supplementary Fig. 1). In addition, the CTCF binding profile in the HeLa-S3 cell line from ChIP-exo experiment was further used to evaluate the prediction performance.

**Fig. 1:**
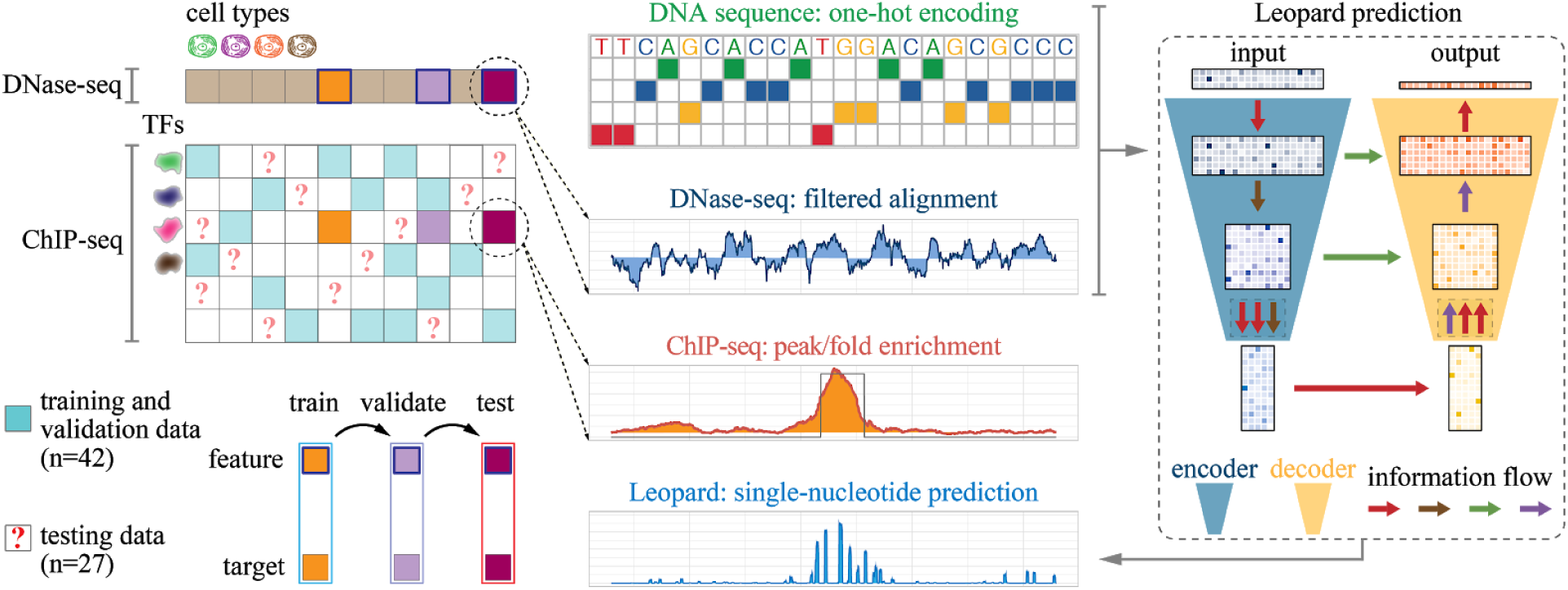
Schematic illustration of Leopard workflow. This study aims to decode the high-resolution transcription factor binding landscapes (refined ChIP-seq signals) based on chromatin accessibility (DNase-seq signals) in a cross-cell type fashion. A total of 42 ChIP-seq experimental results from the ENCODE project were used to train and validate models, whereas the other 27 results were used to test the performance of our method, covering 12 transcription factors. The DNase-seq signals of filtered alignments and one-hot encoded DNA sequences were used as inputs for a deep convolutional neural network model. In contrast to the relatively low-resolution ChIP-seq binding peaks, Leopard generates the cell type-specific transcription factor binding profiles at single-nucleotide resolution.

### The architecture of Leopard captures determinants of TF binding at multiple ranges and resolutions

Leopard was designed to extract information from multiple ranges and resolutions for predicting TF binding locations. Leopard has a deep convolutional neural network architecture, which accepts 6-by-10240 matrices as inputs (Fig. 2a). The 6 channels in the first dimension are signals from (1) DNase-seq, (2) ΔDNase-seq, (3-6) one-hot encoded DNA sequences. The columns in the second dimension correspond to the input length of 10240 successive genomic positions. The Leopard architecture has two components, the encoder and the decoder. The basic building unit of the encoder is the convolution-convolution-pooling (ccp) block, which contains two convolutional layers and one max-pooling layer. In each convolutional layer, the 1D convolution operation is applied to the data along the second dimension (Fig. 2b). By scanning across all 10240 positions, the convolutional layer can capture the upstream and downstream information and key determinants of TF binding will trigger an activation, such as a motif match and open chromatin. Of note, the users do not need to specify the TF motifs since the convolution operators can automatically learn and recognize the regulatory motifs and neighboring DNA sequences. Meanwhile, each max-pooling layer reduces the input length by half, allowing the subsequent convolutional layer to capture the information spanning longer genomic positions. For example, 5 successive points of the input matrix only cover 5 positions whereas 5 successive points at the end of the encoder cover 5*2^5^ = 1600 genomic positions. A total of 5 ccp blocks were used to gradually reduce the input length from 10240 to 320 and increase the number of channels from 6 to 109, compensating the loss of resolution along the genomic dimension. In contrast to the ccp blocks in the encoder, the counterpart upscaling-convolution-convolution (ucc) blocks were used in the decoder. They gradually increase the length but decrease the number of channels. In addition, the concatenation operation transfer the information from the encoder to decoder at each resolution (horizontal green arrows in Fig. 2a). The final output is a 1-by-10240 array, corresponding to the prediction of TF binding signals at each input position at single-base resolution. This neural network architecture effectively captures the long-range and short-range information at multiple scales.

**Fig. 2:**
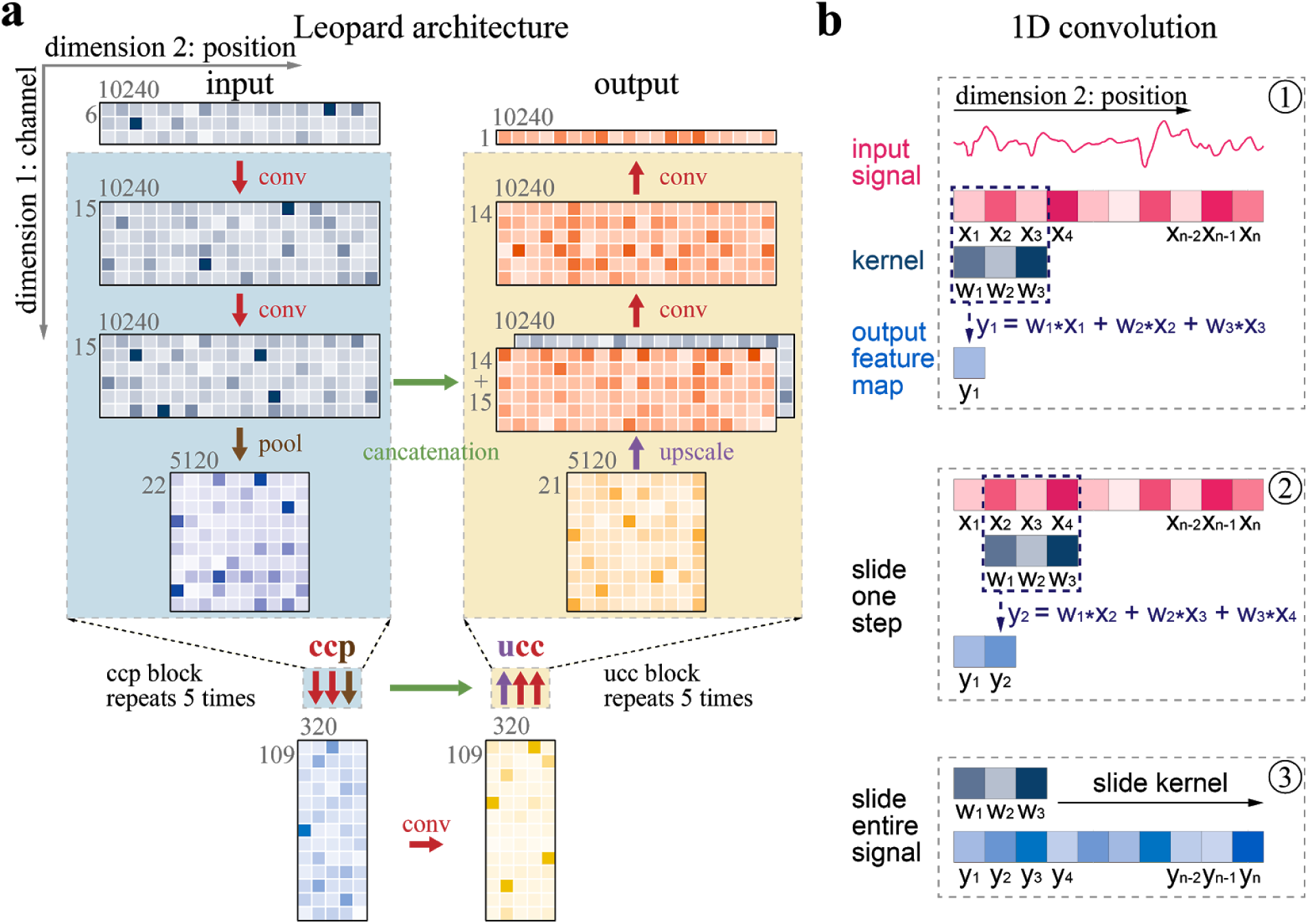
The architecture design of Leopard for single-nucleotide prediction of transcription factor binding. **a**, Leopard accepts 2 dimensional matrices as inputs, where the first dimension represents 6 channels and the second dimension represents 10240 genomic positions. The 6 channels correspond to (1) DNase-seq filtered alignment signals, (2) ΔDNase-seq signals that subtracting the average DNase-seq signal from all cell types under consideration, and (3-6) one-hot encoded DNA sequences. The 10240 genomic positions correspond to randomly sampled consecutive segments in human genome. Leopard has two components: the encoder (blue) and the decoder (yellow). The encoder contains five convolution-convolution-pooling (ccp) blocks that gradually increase dimension 1 and reduce dimension 2. The decoder has five opposite operating units, upscaling-convolution-convolution (ucc) blocks that gradually increase dimension 2 to the same length as the input. This architecture allows for generating outputs for multiple positions simultaneously, substantially boosting the predicting speed. Meanwhile, the concatenation operations (horizontal green arrows) connect the encoder with the decoder, preventing information decay in deep neural networks. **b**, the 1 dimensional (1D) convolution operator calculates the inner product between the kernel (w1, w2, w3) and the input signal (x1, x2, x3), resulting in one feature map value (y1) in step 1. Then the kernel slides along the entire input signal (step 2 and 3) and generates the output feature map vector, which has the same size in dimension 2 as the input.

### Leopard accurately identifies cell type-specific TF binding profiles at single-nucleotide resolution

We first compared the Leopard prediction profiles with the refined ChIP-seq conservative peaks in the 27 testing TF-cell type pairs. For each TF, we trained and validated our model in a subset of training cell types, then made predictions on the other subset of testing cell types. The complete train-validate-test partition is shown in Supplementary Fig. 1. Both the area under receiver operating characteristic curve (AUROC) and the area under precision-recall curve (AUPRC) were calculated and compared (Fig. 3a-b; Supplementary Fig. 2; Supplementary Table 1). Leopard accurately identified TF binding profiles for the 12 TFs in this study, with a median prediction AUROC of 0.994. In 19 out of 27 (70%) testing TF-cell type pairs, Leopard achieved high AUROCs above 0.990. The corresponding precision recall curves and AUPRCs are shown in Supplementary Fig. 2. Of note, identifying TF binding sites is an extremely difficult class imbalance problem - only 0.11% of all genomic positions were bound by TFs. The AUPRC baseline of random prediction was therefore very low around 0.0011 (the dashed line in Supplementary Fig. 2). Compared with the random prediction baseline, Leopard had more than hundredfold improvement. The AUROC and AUPRC results demonstrated the high prediction accuracy of our method.

**Fig. 3:**
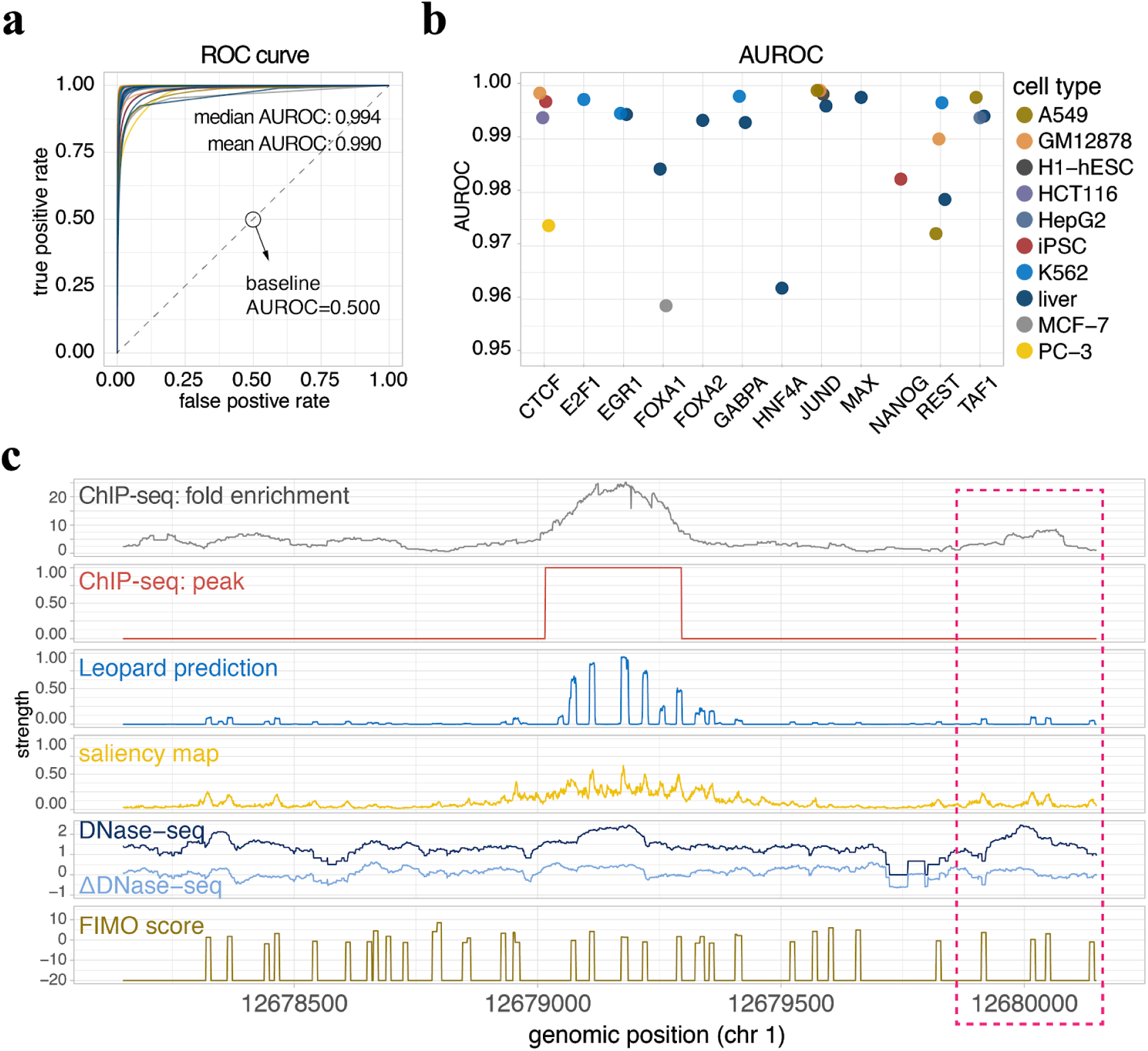
Leopard identifies cell type-specific transcription binding events at single-nucleotide resolution. **a**, The receiver operating characteristic (ROC) curves of Leopard predictions on 27 held-out testing TF-cell type pairs. The baseline score of random prediction is 0.500 shown as the dashed line along the diagonal. **b**, Leopard achieves high area under receiver operating characteristic curves (AUROCs) for 12 transcription factors in 10 testing cell types. Each dot represents the overall AUROC calculated from the testing Chr1, Chr8, and Chr21. Different colors represent different cell types. **c**, An example 2000bp segment is shown to demonstrate the prediction results. This segment contains signals between genomic positions 12,678,147 and 12,680,147 of Chr 1 from the JUND binding profile in the A549 cell line. The top two rows are the original ChIP-seq fold enrichment and conservative peaks generated through the standard ENCODE analysis pipeline. The ChIP-seq broad binding peak can not locate the precise binding site. Leopard generates single-nucleotide predictions and precisely provides the potential binding sites. The saliency map indicates positions contributing to the predictions. The corresponding DNase-seq and ΔDNase-seq signals, and the sequence-based motif scan scores using FIMO are also shown here for comparison. Of note, the region in the pink rectangle also has open chromatin (high DNase-seq signals) and binding motifs (high FIMO scores), but no binding events were observed from the ChIP-seq experiment. Leopard is able to detect these non-binding locations - no prediction peaks in this region.

To visualize the Leopard prediction results, we demonstrate a 2000bp segment of JUND binding profile in A549 cell line as an example (Fig. 3c). The raw ChIP-seq data were processed through the standard ENCODE analysis pipeline, resulting in a broad region of putative binding sites between locations 12,678,147 and 12,680,147 in chromosome (Chr) 1, in terms of both the fold enrichment and conservative peaks that pass the 5% IDR threshold (the top two rows in Fig. 3c). These relatively low-resolution ChIP-seq profiles can not provide the precise binding locations. In contrast, the predictions from Leopard clearly depicted the exact binding locations, overlapping with the ChIP-seq peaks (the third row in Fig. 3c). The saliency map was also calculated, indicating positions that contributed to the predictions [25]. For comparison, we also aligned the DNase-seq signal, the ΔDNase-seq signal, and FIMO motif scanning score. As we expected, without the cell type-specific information on chromatin accessibility, the sequence-based motif scanning approach generates many false positive binding sites (peaks in the bottom row in Fig. 3c). On the other hand, the DNase-seq and ΔDNase-seq signals indicated the open chromatin regions, which were prerequisite for TF binding except for pioneering TFs. In general, a genomic site with the “open chromatin” status and a high motif match score is more likely but not necessary to be a binding site. Leopard is able to distinguish true binding events from false ones. For instance, the region around location 12,680,000 in the pink rectangle in Fig. 3c has high DNase-seq signals and multiple motif matches. However, no binding events were observed within this region based on the ChIP-seq experiment. Leopard correctly predicted no binding peaks within this region.

We further compared the above 2000bp genomic region across four cell types (A549, GM12878, H1-hESC, and liver) and observed distinct TF binding patterns across cell types (Fig. 3c; Supplementary Fig. 3). In A549 (Fig. 3c), H1-hESC (Supplementary Fig. 3b), and liver (Supplementary Fig. 3c) cell types, JUND had a similar binding peak in this region and Leopard correctly predicted these binding events. In contrast, JUND did not bind to this region in GM12878 (Supplementary Fig. 3a) cell line, potentially due to the relatively restricted chromatin accessibility in this specific cell line. Leopard successfully distinguished GM12878 from the other three cell types, since DNase-seq and ΔDNase-seq signals were the cell type-specific input features for Leopard. In contrast, computational methods only based on sequences without DNase-seq features will not identify the differences across cell types. These results demonstrate the great advantage of Leopard over sequence-based methods for predicting TF binding sites.

### Benchmarking Leopard prediction against the single-base ChIP-exo experiment

To further evaluate the prediction performance of Leopard at single-base resolution, we compared our predictions with CTCF binding peaks generated from ChIP-exo experiment. The receiver operating characteristic (ROC) curves and precision recall (PR) curves on each individual chromosome (Chr 1-22, X) and overall across human genome are shown in Fig. 4a-b (Supplementary Table 2). The over AUROC of 0.990 is very high and the AUPRC score of 0.297 indicating that Leopard achieved more than 500-fold improvement over the AUPRC baseline of 0.00065 (the dashed line in Fig. 4b). Of note, Leopard was only trained on the low-resolution ChIP-seq data, yet it was able to identify the high-resolution ChIP-exo binding peaks. In Fig. 4c, we demonstrated an example prediction of the CTCF binding region in the HeLa-S3 cell line. For comparison, the corresponding ChIP-seq, ChIP-exo, DNase-seq, and motif-based scanning signals were also aligned and shown in Fig. 4c. As we expected, the ChIP-exo experiment has a much narrower peak (the third row in Fig. 4c) than the fold enrichment or the conservative peak from the ChIP-seq experiment. Leopard precisely predicted the binding sites, corresponding well to the ChIP-exo peaks. In summary, Leopard can identify the single-base TF binding profiles based on chromatin accessibility and DNA sequence.

**Fig. 4:**
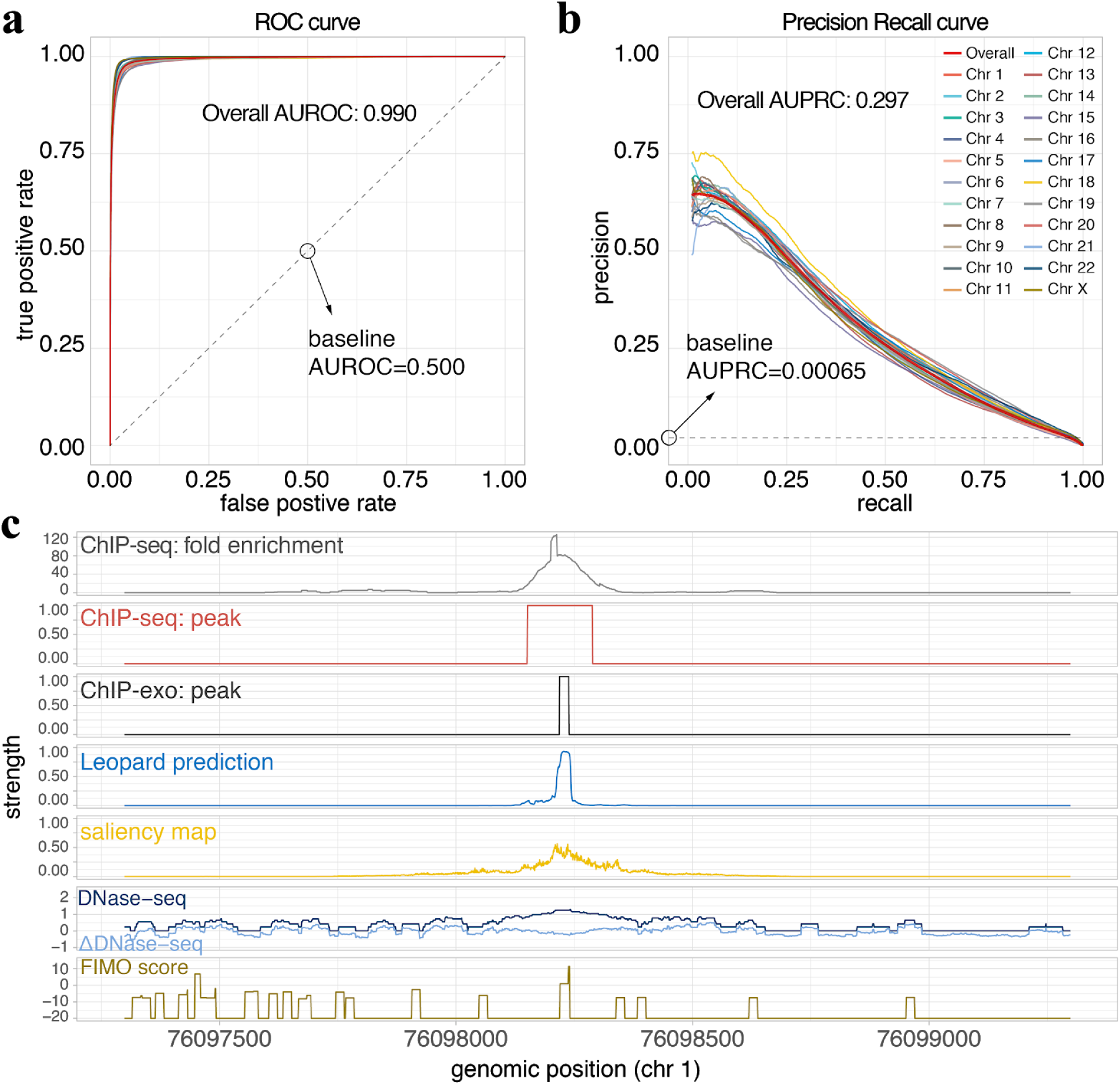
Evaluating Leopard predictions based on TF binding profiles from the ChIP-exo experiment. **a**, The receiver operating characteristic (ROC) curve and **b**, precision recall (PR) curve between Leopard prediction and peaks from *in vivo* ChIP-exo experiment of the CTCF binding profile in HeLa-S3 cell line. The ROC and PR curves were evaluated and drawn on each individual chromosome and the overall genome-wide performance of all chromosomes (Chr 1-22 and X). The baseline of random predictions is shown as dashed lines. **c**, An example 2000bp segment is shown to demonstrate Leopard prediction results. This segment contains signals between genomic positions 76,097,298 and 76,099,298 of Chr 1 from the CTCF binding profile in the HeLa-S3 cell line. The top two rows are the original ChIP-seq signals from the fold coverage change and conservative peak profiles generated through the standard ENCODE analysis pipeline. The third row is the ChIP-exo signal precisely describing the binding site. Leopard generates single-nucleotide predictions and the prediction corresponds well with the ChIP-exo peak. The saliency map indicates positions contributing to the predictions. The corresponding DNase-seq and ΔDNase-seq signals, and the sequence-based motif scan scores using FIMO are also shown here for comparison.

### Leopard substantially outperforms state-of-the-art methods for predicting TF binding site despite evaluated at the low 200bp resolution

We further compared state-of-the-art methods for TF binding site prediction, which were the top performing methods named Anchor and FactorNet in the ENCODE-DREAM *in vivo* Transcription Factor Binding Site Prediction Challenge. Anchor is based on a classical treed-based machine learning model, XGBoost, to predict TF binding sites through sophisticated feature engineering [22], whereas FactorNet is based on a many-to-one neural network model to address this problem. Notebly, these methods only provided predictions at the low 200bp resolution and it is not a fair comparison for Leopard. Nevertheless, we adapted Leopard architecture to generate predictions at 200bp resolution, which was called Leopard-200bp (Supplementary Fig. 4). We benchmarked Leopard-200bp against Anchor and FactorNet on the same training and testing datasets in the ENCODE-DREAM Challenge (Supplementary Fig. 5). The performance was evaluated in a cross-cell type and cross-chromosome fashion (see details in Methods). Overall, Leopard achieved higher prediction AUPRCs than Anchor and FactorNet (Fig. 5a-b; Supplementary Table 3). The mean and median AUPRCs on the right clearly demonstrate the improvement of Leopard predictions in the 13 testing TF-cell type pairs. The percentage improvement over Anchor and FactorNet was further calculated. Leopard substantially improved the median prediction AUPRC by 19% and 27% over Anchor and FactorNet, respectively. We further performed pairwise statistical comparison of these methods (Supplementary Fig. 6). For each testing TF-cell type pair, we randomly sampled 100 segments with length of 100kbp then calculated 100 prediction AUPRCs. The paired Wilcoxson signed rank test was performed. Leopard significantly outperformed Anchor and FactorNet in the overwhelming majority of testing TF-cell type pairs.

**Fig. 5:**
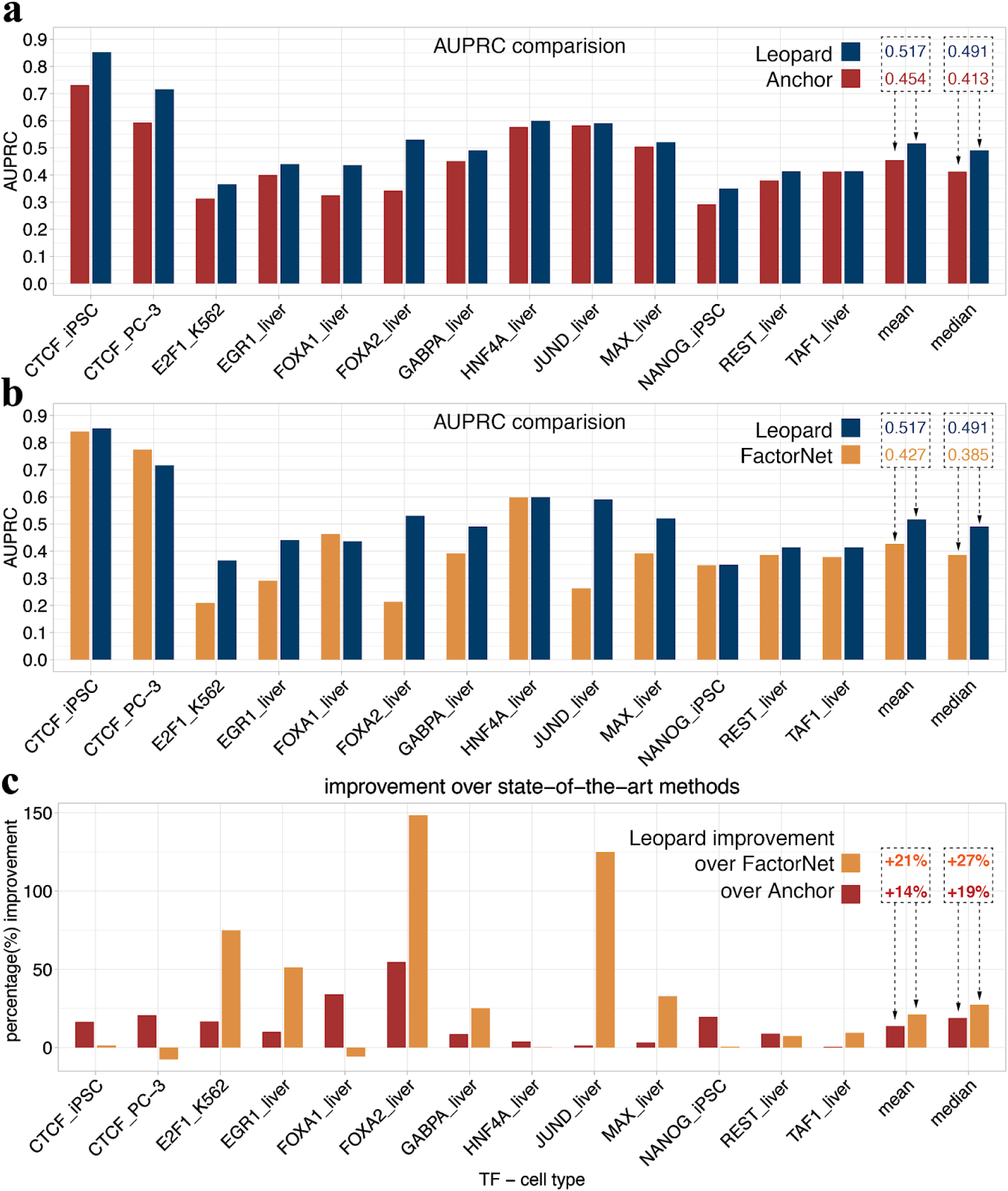
Leopard substantially improved transcription factor binding site prediction over state-of-the-art methods. Leopard was benchmarked with **a**, Anchor (the top performing classical tree-based model) and **b**, FactorNet (the top performing neural network model) in the ENCODE-DREAM *in vivo* Transcription Factor Binding Site Prediction Challenge. The same challenge training and testing datasets were used to train Leopard model and evaluate performance. Overall, Leopard achieved higher prediction AUPRCs than Anchor and FactorNet. The mean and median AUPRCs on the right clearly demonstrate the advantage of Leopard predictions in the 13 testing TF-cell type pairs. **c**, The percentage improvement over Anchor and FactorNet was calculated. Leopard substantially improved the median prediction AUPRC by 19% and 27% over Anchor and FactorNet, respectively.

In addition to higher prediction resolution and accuracy, Leopard has a great speed advantage over previous methods. Previous neural network approaches were typically based on the many-to-one architecture [18–21,23], in which genomic segments containing hundreds or thousands of base pairs were used as inputs to predict only a single label at a time. Considering the fact that the human genome contains more than 3 billion base pairs, it requires considerably long runtime for the genome-wide predictions using many-to-one models. In contrast, Leopard is based on a many-to-many neural network architecture, in which the base-wise label for every nucleotide in the input genomic segment (10240bp) are generated simultaneously. This unique architecture tremendously boosts the prediction speed. To benchmark the runtimes of the many-to-one and many-to-many neural network structures, we modified Leopard and created a many-to-one version that accepted 10240bp as inputs and generated only one prediction value (Supplementary Fig. 7). For comparison, we also tested the runtime of Anchor, representing the speed of tree-based XGBoost model. Leopard can finish prediction within 4.70, 2.80, and 0.95 minutes for predicting the single-nucleotide HNF4A binding sites on Chr1, Chr8, and Chr21 respectively (Fig. 6a). Yet Anchor and the many-to-one neural network requires much longer runtimes on the scales of hours. Meanwhile, Leopard is flexible with both graphics processing unit (GPU) and central processing unit (CPU) settings. When running Leopard on CPU, the prediction runtimes remain acceptable on the scale of minutes (Fig. 6b). When making genome-wide predictions at single-nucleotide resolution, Leopard has a great advantage of speed - it only requires less than one hour on GPU or four hours on CPU. In contrast, it requires 583.6 hours (>24 days) for the many-to-one neural network model and 1220.8 hours (>50 days) for Anchor (Fig. 6c). Therefore, Leopard not only outperformed but also enabled hundred-fold to thousand-fold speedup over the common many-to-one neural network and the Anchor model, respectively.

**Fig. 6:**
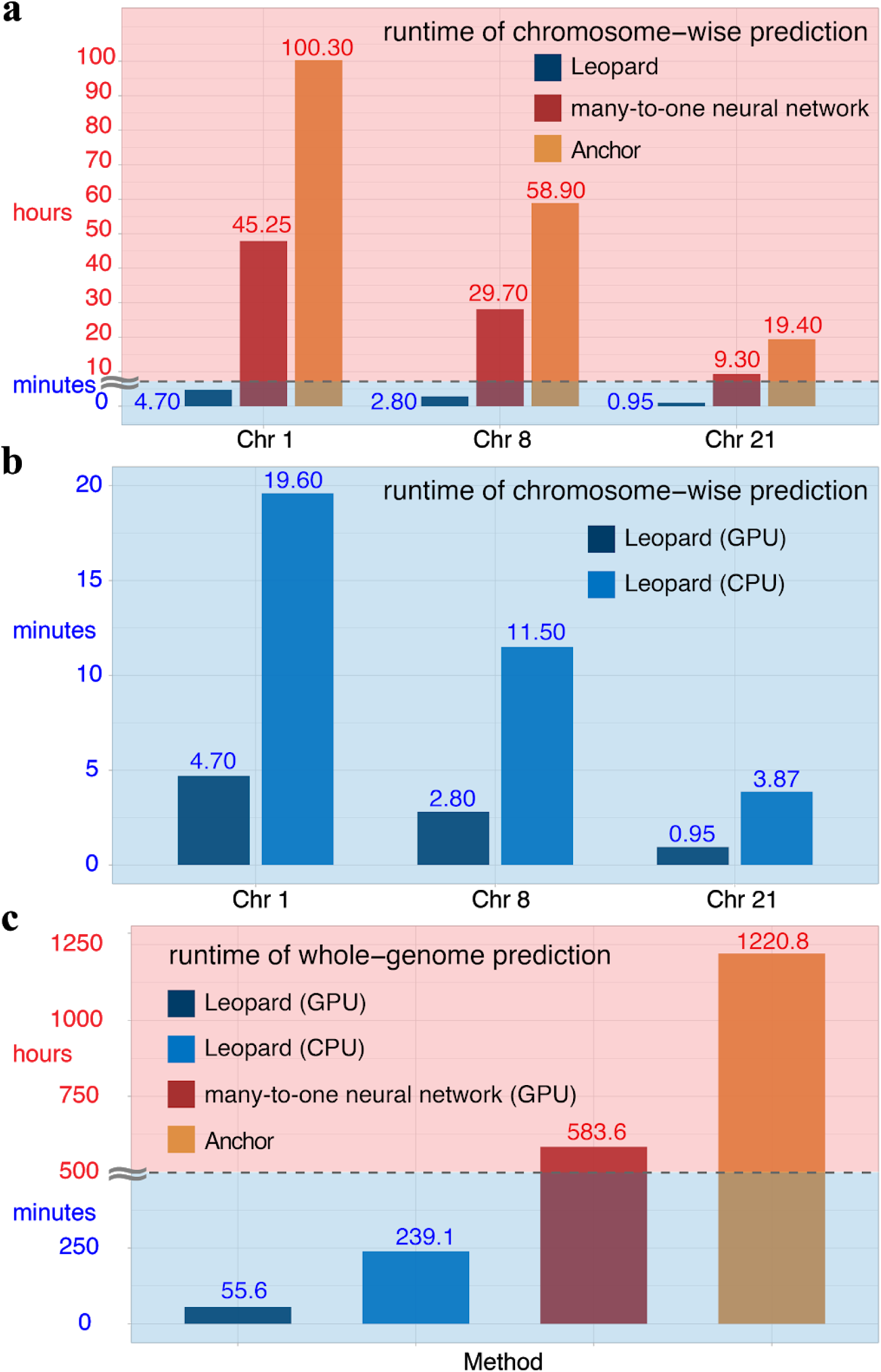
The runtimes of chromosome-wise and whole-genome predictions of Leopard and other state-of-the-art methods. The runtime was tested for predicting the HNF4A binding profiles in the liver cell using different methods. **a**, The runtime was tested for chromosome-wise predictions on Chr 1, Chr 8 and Chr 21. Notably, the runtimes of bars within the blue background are in the unit of minute, whereas the runtimes in the red background are in the unit of hour. For example on Chr 8, it only takes 2.8 minutes for Leopard to finish predictions whereas it requires 29.70 and 58.90 hours for the many-to-one neural network model and the Anchor method using XGBoost. Leopard is therefore estimated to be 600 and 1200 times faster than the many-to-one model and Anchor on average. **b**, Leopard is flexible with both GPU and CPU settings. When running Leopard on CPU, the prediction runtimes are acceptable on the scale of minutes. The runtime of CPU was tested on a standard computer with eight CPU cores. **c**, When making genome-wide predictions at single-nucleotide resolution, Leopard has a great advantage of speed - it only requires 55.6 minutes on GPU or 239.1 minutes on CPU. In contrast, it requires 583.6 hours (>24 days) for the many-to-one neural network model and 1220.8 hours (>50 days) for Anchor.

## Discussion

Many studies have proposed computational models for predicting the sequence specificities of DNA-binding proteins and variant effects *ab initio* from DNA sequences [18,19,26,27]. These approaches have advanced our understanding of the transcriptional regulation. However, in the new era of precision medicine, the context-dependent functional genomic and epigenomic landscapes across different cell types, tissues, and patients may not be completely encoded in the DNA sequence space. These genomic and epigenomic profiles provide crucial information for disease mechanism investigation and future treatment discovery. In this work, we develop a robust, scalable, and fast software, Leopard, to identify cell type-specific TF binding sites at the single-nucleotide level, which is a key challenge in computationally decoding functional elements of the human genome. Leopard leverages the cutting-edge deep learning framework for pixel-level image segmentation and large-scale data from the ENCODE project, achieving high cross-cell type prediction accuracy.

Unlike classical machine learning approaches that require complicated feature crafting and engineering [22,28,29], neural network models automatically learn the informative features and factors contributing to TF binding, largely increasing the flexibility and scalability [30,31]. Moreover, the limited hard drive size and computer memory usually restrict the complete search of the feature space in classical machine learning approaches. For example, both Anchor [22] and the method reported by J-team [28] calculated bin-level aggregate statistical features (maximum, mean, minimum, and other statistics) to represent the major signals instead of using the complete signals for the sake of computational efficiency. In contrast, Leopard accepts the complete DNase-seq signals of the input segments covering 10240 positions without information loss, which may explain its higher predictive performance. In terms of network structure tuning, we adapted the standard U-Net structure into the 1D version, which demonstrated improved performance over the state of the art. In fact, recent work in the field of neural architecture search indicates that neural networks based on randomly wired graphs achieved competitive performance to manually designed architectures on image classification [32]. It would be interesting to see the future application of these models to bioinformatics research.

Recent advancement in deep learning studies of genomic data has generated novel interpretations and biological hypotheses [16,17]. Although the many-to-one neural network (with various convolutional, recurrent, and fully connected layers) has been actively explored, the many-to-many neural network remains to be developed. A major advantage of the many-to-many structure is the ultrafast prediction speed, which will accelerate the genomic research on the large genome-wide scale at single-nucleotide resolution. Here we demonstrate its very first application to identify TF binding events. We envision that our analysis framework and model can be flexibly adapted to investigate many other genomics modeling tasks in the future, including predicting various epigenetic modifications in different cell types.

## Conclusions

In this study, we present a novel deep neural network approach, Leopard, for decoding cell type-specific TF binding landscapes at single-nucleotide resolution. Leopard identified high-resolution ChIP-exo TF binding locations although it was only trained on low-resolution ChIP-seq signals. Meanwhile, Leopard leverages a many-to-many deep learning framework to generate predictions for multiple genomic positions simultaneously, allowing for hundred-fold to thousand-fold speedup compared with current many-to-one deep learning and classical machine learning models.

## Methods

### Data collection and preprocessing

In this study, we used the data from the ENCODE project, which consists of 69 ChIP-seq experiments and 13 DNase-seq experiments covering 12 TFs (CTCF, E2F1, EGR1, FOXA1, FOXA2, GABPA, HNF4A, JUND, MAX, NANOG, REST, and TAF1) in 13 cell types (A549, GM12878, H1-hESC, HCT116, HeLa-S3, HepG2, iPSC, IMR-90, K562, liver, MCF-7, Panc1, PC-3). A subset of 27 ChIP-seq experiments were held out as the evaluation testing set and the remaining ChIP-seq data were used for model training (Supplementary Fig. 1). The ChIP-seq data were processed following the standard ENCODE analysis pipeline. For each ChIP-seq experiment, the raw sequencing reads were mapped to hg19 reference genome using BWA aligner [33]. The SPP peak caller was used to call up to 300,000 peaks in each ChIP-seq replicate over the control using a relaxed false discovery rate of 0.9 [34]. Then the irreproducible discovery rate (IDR) method was used to find the reproducible and conservative peaks with the IDR threshold of 5% [35]. The fold enrichment signals were generated using the MACSv2 peak caller [36]. To generate the high-resolution training labels, we further refined the ChIP-seq conservative peaks by overlapping them with the motif scanning hits. We scanned the hg19 reference genome using FIMO [24] based on the TF motifs from Hocomoco11 [8]. A relaxed cutoff of p=0.01 was used to locate all the potential binding sites. In addition, we downloaded the ChIP-exo data [5] and generated peaks using the MACE analysis pipeline [37].

When comparing our method with the top-ranking methods in the ENCODE-DREAM challenge, we used the same dataset provided by the challenge consisting of 43 and 13 ChIP-seq data for model training and held-out testing, respectively (Supplementary Fig. 5). Of note, in the challenge, the resolution of predictions was only 200bp. For each 200bp interval at 50bp sliding window steps, a gold standard binary label “Bound” or “Unbound” was assigned. The AUPRC between predictions and gold standard labels was used as the scoring metric to compare different models, which was also used in the ENCODE-DREAM challenge. For this low-resolution prediction task, we created a Leopard-200bp model (Supplementary Fig. 4), which directly generate predictions at the 200bp resolution. In addition to the one-hot encoded sequences and DNase-seq features, we also added the orange feature (the 5’ tag count within each 200bp bin) [38], which was also named as frequency/Δfrequency feature in Anchor [22].

For the 13 DNase-seq data, we used raw filtered alignment files without peak calling. Signals from multiple technical and biological replicates of the same cell type were summed and combined. To reduce the potential cell-specific cleavage and sequencing biases, we performed the quantile normalization following the same pipeline as described in Anchor [22].

### AUROC and AUPRC

The area under receiver operating characteristic curve (AUROC) and the area under precision-recall curve (AUPRC) between prediction and gold standard were used to evaluate the prediction performance. Since the predictions are continuous values between 0 and 1, a series of cutoff values, [0, 0.001, 0.002, …, 0.998, 0.999, 1.000], are used to binarize the predictions. At each cutoff, the True Positive Rate (TPR) and the False Positive Rate (FPR) are defined as

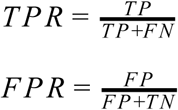

where TP is True Positive, FN is False Negative, FP is False Positive, and TN is True Negative. Similarly, the Precision and Recall are defined as

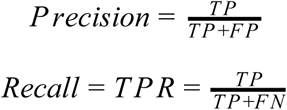

These values were calculated at each cutoff, forming the receiver operating characteristic curve and precision-recall curve. The area under the curve therefore reflects the prediction performance of a model.

### Overall AUPRC and AUROC

The overall AUPRC, or the gross AUPRC, is defined as

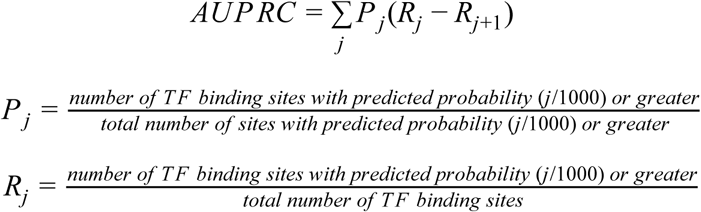

where the Precision (*P* _*j*_) and Recall (*R*_*j*_) were calculated at each cutoff j and j = 0, 0.001, 0.002, …, 0.998, 0.999, 1. When multiple chromosomes are under consideration, this overall AUPRC is similar to the “weighted AUPRC”, which is different from simply averaging the AUPRC score of all chromosome [30]. This is because the overall AUPRC considers the length of each chromosome and longer chromosomes contribute more to the overall AUPRC, resulting in a more accurate evaluation of the performance. The overall AUROC is defined in a similar way as the overall AUPRC.

### Convolutional neural network architecture

The architecture of Leopard was adapted from the image segmentation neural network model, U-Net, which generates pixel-wise labels for every pixel in the input 2D image [39]. Similarly, Leopard generates base-wise labels for every nucleotide in the input genomic segment (10240bp) simultaneously. In each convolutional layer, the kernel size of 7 was used. In each pooling layer, max pooling was used. Each convolution operation was followed by a nonlinear activation, Rectified Linear Unit (ReLU), which is defined as

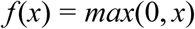

where x is the input and f(x) is the output. Only positive values active a neuron and ReLU allows for effective training of neural networks compared to other complex activation functions. In addition, batch normalization was used after each convolutional layer. In the final layer, we used the sigmoid activation unit to restrict the prediction value between 0 and 1. The sigmoid activation is defined as

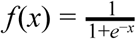

where x is the input and f(x) is the output.

### Cross-cell type and cross-chromosome training, validation, and testing

We used a “crisscross” training and validation strategy to build models and avoid overfitting [22]. For each TF, we first collected all the available training cell types. Then a pair of cell types was selected, one for model training and the other for model validation. Meanwhile, the 23 chromosomes (Chr 1-22 and X) were also partitioned into the training, validation, and testing sets. Chr 1, Chr 8, and Chr21 were fixed throughout this study as the testing chromosome set,whereas the remaining 20 chromosomes were randomly partitioned into the training and validation sets.

During model training, we defined an epoch as 100,000 segments or samples randomly selected from the training chromosome set in the training cell type. Each time after the model was trained on one epoch of training samples, another epoch of validation samples were randomly selected from the validation chromosome set in the validation cell type to calculate the prediction losses, monitor the training progress, and avoid overfitting. The Adam optimizer was used. Each neural network model was first trained for 5 epochs with the learning rate of 1e-3 and then trained for 15 epochs with the learning rate of 1e-4 till the loss converged.

### Training Losses

The cross entropy loss, was used for model training, which is defined as

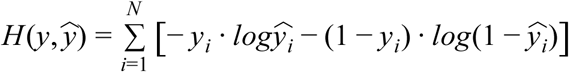

where *y*_*i*_ is the gold standard label of TF binding = 1 or non-binding = 0 at genomic position i, 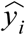 is the prediction value at position i, N=10240 is the total number of base pairs in each segment, *y* is the vector of the gold standard labels and 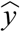 is the vector of predictions. Ideally, an “AUPRC loss” should be used for optimizing the AUPRC. However, the AUPRC function is not mathematically differentiable, which is required by the back-propagation algorithm during neural network model training. We therefore used the cross-entropy loss to approximate the “AUPRC loss”.

## Declarations

## Data availability

The ChIP-seq data were downloaded from the ENCODE-DREAM challenge website: https://www.synapse.org/#!Synapse:syn6181337 (conservative peaks) https://www.synapse.org/#!Synapse:syn6181334 (fold enrichment) and the ENCODE project website: https://www.encodeproject.org/ (The accession numbers are provided in Supplementary Table 4.)

The DNase-seq data were downloaded from the ENCODE-DREAM challenge website: https://www.synapse.org/#!Synapse:syn6176232

## Software availability

The source code of our Leopard software is available on GitHub: https://github.com/GuanLab/Leopard

## Competing interests

YG receives personal payment from Eli Lilly and Company, Genentech Inc, F. Hoffmann-La Roche AG, and Cleerly Inc; holds equity shares at Cleerly Inc and Ann Arbor Algorithms Inc; receives research support from Merck KGaA as research contracts and Ryss Tech as unrestricted donations.

## Funding

This work is supported by NSF-US14-PAF07599 (#1452656; CAREER: On-line Service for Predicting Protein Phosphorylation Dynamics Under Unseen Perturbations NSF), R35-GM133346-01 (Machine Learning for Drug Response Prediction), AWD007950 (Digital Biomarkers in Voices for Parkinson’s Disease American Parkinson’s Disease Association), University of Michigan O’Brien Kidney Translational Core Center, and Michael J. Fox Foundation #17373 to YG. This work is also supported by 19AMTG34850176 (American Heart Association and Amazon Web Services3.0 Data Grant Portfolio: Artificial Intelligence and Machine Learning Training Grants) to HL. We thank the GPU donation from Nvidia and the AWS donation from Amazon.

## Authors’ contributions

YG and HL conceived and designed this project. HL performed the experiments, prepared the figures and wrote the manuscript. All authors contributed to the writing of the manuscript and approved the final manuscript.

