## Supplemental Information for "Leopard: fast decoding cell type-specific transcription factor binding landscape at single-nucleotide resolution"

Contents

The system configuration to test runtimes …….………………….………..…………………. 2

Supplementary Fig. 1 ……………………………………………………………………….. 3
Supplementary Fig. 2 ……………………………………………………………………….. 4
Supplementary Fig. 3 ……………………………………………………………………….. 5
Supplementary Fig. 4 ……………………………………………………………………….. 7
Supplementary Fig. 5 ……………………………………………………………………….. 8
Supplementary Fig. 6 ……………………………………………………………………….. 9
Supplementary Fig. 7 …………………………………………………...………………….. 10
Supplementary Table 1 ……………………………………………………………………..… 11
Supplementary Table 2 ……………………………………………………………………... 12
Supplementary Table 3 ……………………………………………………………………..… 13
Supplementary Table 4 ……………………………………………………………………... 13

**The system configuration to test runtimes**

**GPU**

NVIDIA GeForce GTX TITAN X 12GB

**CPU**

Architecture: x86_64

CPU op-mode(s): 32-bit, 64-bit

Byte Order: Little Endian

CPU(s): 8

On-line CPU(s) list: 0-7

Thread(s) per core: 2

Core(s) per socket: 4

Socket(s): 1

NUMA node(s): 1

Vendor ID: GenuineIntel

CPU family: 6

Model: 158

Model name: Intel(R) Core(TM) i7-7700K CPU @ 4.20GHz

Stepping: 9

CPU MHz: 800.061

CPU max MHz: 4500.0000

CPU min MHz: 800.0000

BogoMIPS: 8400.00

Virtualization: VT-x

L1d cache: 32K

L1i cache: 32K

L2 cache: 256K

L3 cache: 8192K

NUMA node0 CPU(s): 0-7

**Memory**

31GB in total

**System**

Linux version 4.15.0-62-generic (buildd@lcy01-amd64-024) (gcc version 7.4.0 (Ubuntu 7.4.0-1ubuntu1~18.04.1)) #69-Ubuntu SMP Wed Sep 4 20:55:53 UTC 2019

### Supplementary Figures

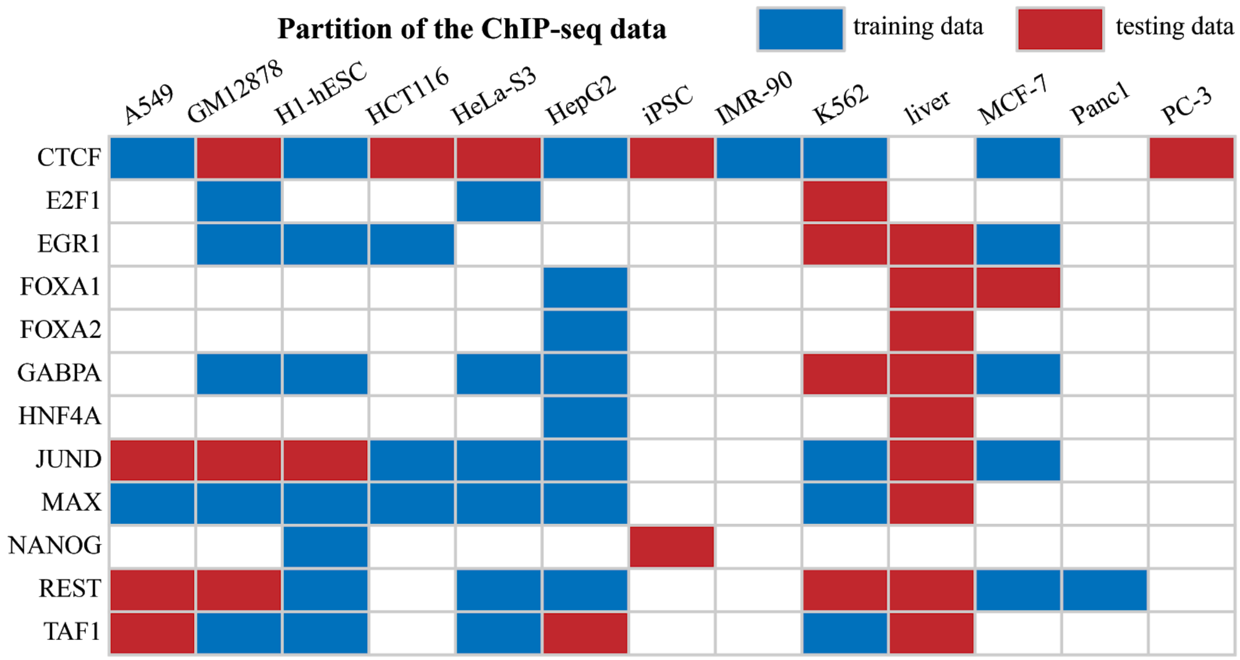


Supplementary Fig. 1: The partition of dataset for model training and testing.

A total of 69 ChIP-seq experiments the ENCODE project were used, covering 12 TFs (CTCF, E2F1, EGR1, FOXA1, FOXA2, GABPA, HNF4A, JUND, MAX, NANOG, REST, and TAF1) in 13 cell types (A549, GM12878, H1-hESC, HCT116, HeLa-S3, HepG2, iPSC, IMR-90, K562, liver, MCF-7, Panc1, PC-3). A subset of 42 ChIP-seq experiments were used for model training (blue blocks) and the remaining 27 were held-out for model testing (red blocks)

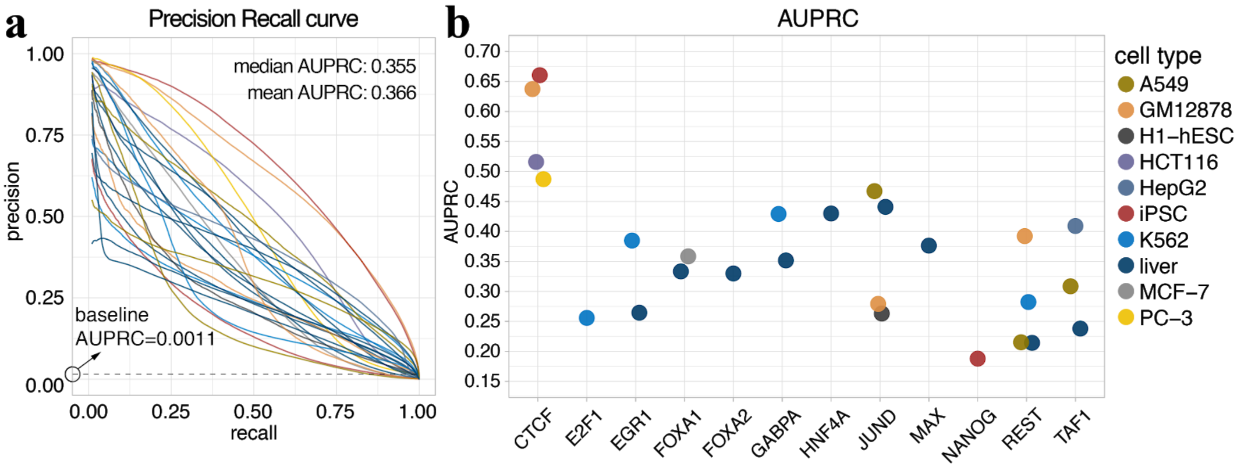


Supplementary Fig. 2: The prediction precision recall curves and the areas under the curves of Leopard predictions evaluated on refined ChIP-seq signals at single-nucleotide resolution.

**a,** The precision recall (PR) curves of 27 testing TF-cell type pairs. The baseline score of random prediction is 0.0011 shown as the dashed line, corresponding to the number of TF binding sites over the total number of base pairs in chromosomes under consideration (Chr1, Chr8, and Chr21). **b,** Leopard achieves high areas under the precision recall curves (AUPRCs) for 12 transcription factors in 10 testing cell types. Each dot represents the overall AUROC calculated from the testing Chr1, Chr8, and Chr21. Different colors represent different cell types.

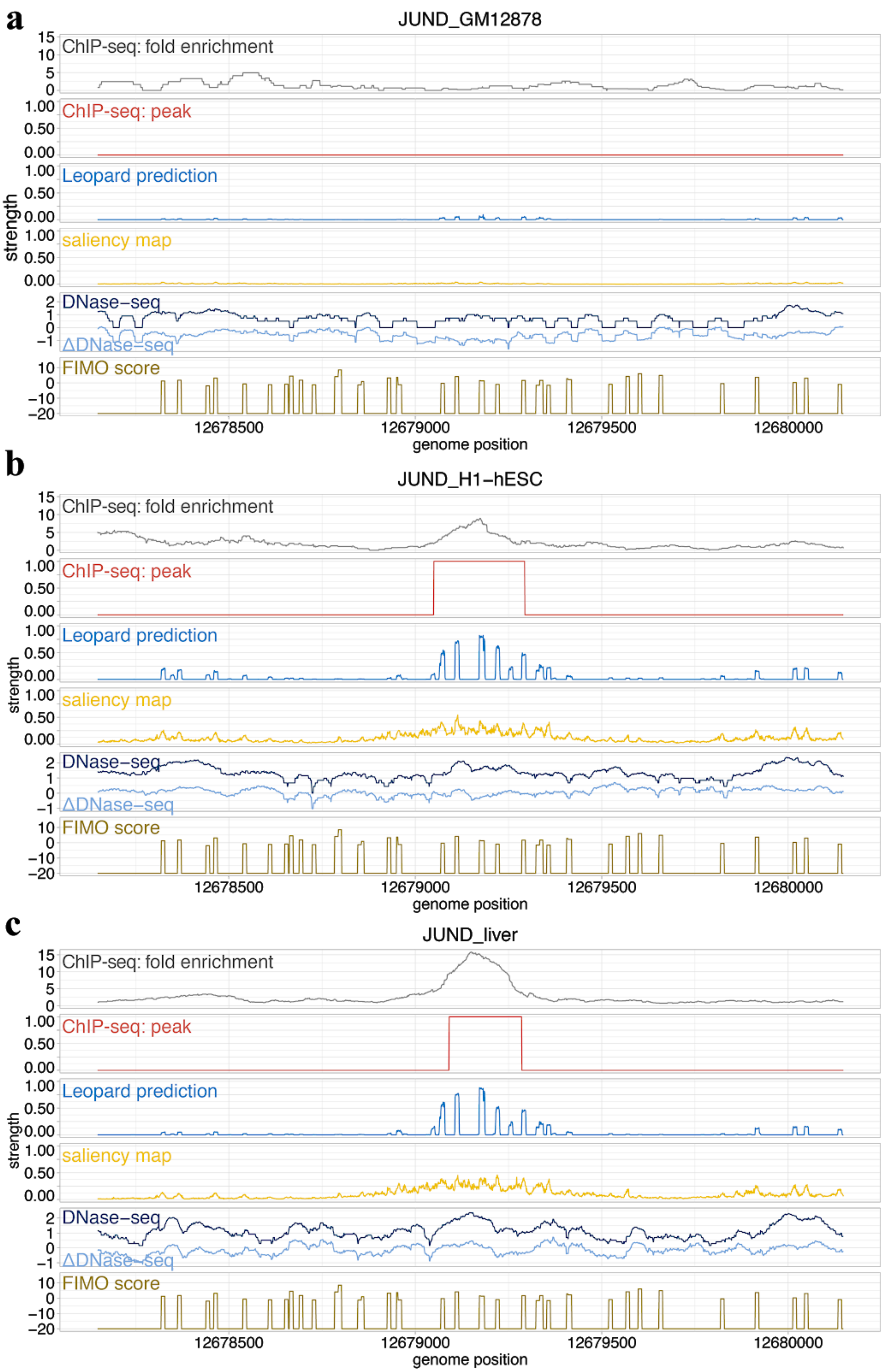
a
JUND_GM12878
b
JUND_H1−hESC
c
JUND_liver
ChIP-seq: fold enrichment
ChIP-seq: peak
Leopard prediction
saliency map
DNase−seq
ΔDNase−seq
FIMO score
strength
genome position
12678500
12679000
12679500
12680000

Supplementary Fig. 3: Leopard distiguishes cell type-specific transcription binding and non-binding events at single-nucleotide resolution.

An example 2000bp segment is shown to demonstrate Leopard prediction results across three cell types: **a,** GM12878 **b,** H1-hESC and **c,** liver. This segment contains successive signals between genomic positions 12,678,147 and 12,680,147 of Chr 1 from the JUND binding profiles. The top two rows are the original ChIP-seq fold enrichment and conservative peaks generated through the standard ENCODE analysis pipeline. The ChIP-seq broad binding peak can not locate the precise binding site. Leopard generates single-nucleotide predictions and precisely provides the potential binding sites in H1-hESC and liver cell types. The saliency map indicates positions contributing to the predictions. The corresponding DNase-seq and ΔDNase-seq signals, and the sequence-based motif scan scores using FIMO are also shown here for comparison. Of note, no JUND binding events were observed by ChIP-seq experiments in GM12878 cell type. The sequence-based models such as FIMO motif scanning will not distinguish this cell type-specific signals. In contrast, Leopard successfully depicted the differences across cell types and made no peak predictions only in GM12878.

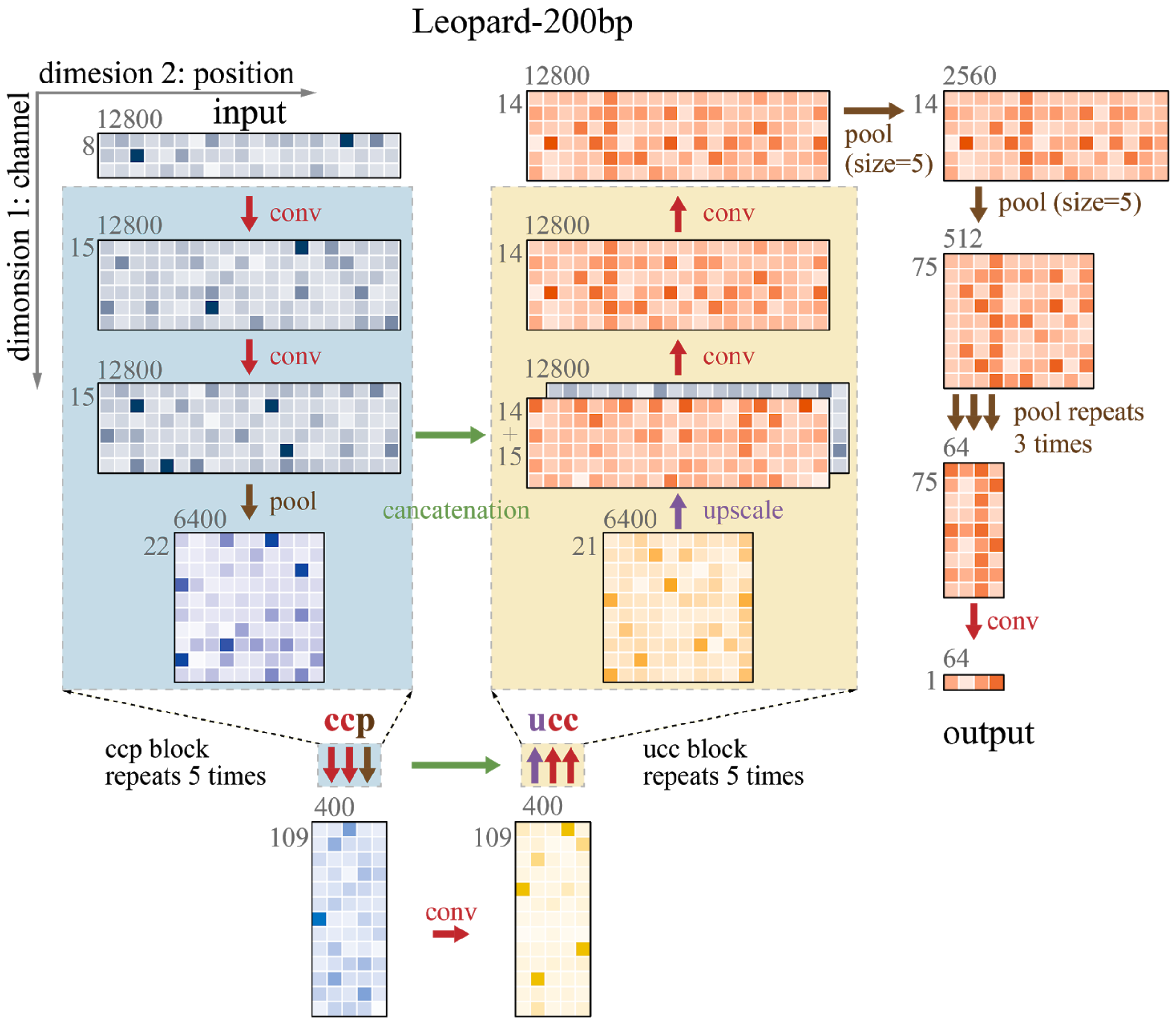


Supplementary Fig. 4: The architecture of Leopard-200bp designed for predicting genome-wide TF binding sites at 200bp resolution.

The encoder (blue) and decoder (yellow) components of Leopard-200bp are similar to those of Leopard, except for the input length was adjusted from 10240 to 12800. We made this adjustion because 12800 (instead of 10240) can be divided by 200 so that the output is a 1-by-64 array corresponding predictions at 200bp resolution. This is accomplished by multiple max-pooling layers (brown arrows on the right) gradually reducing the size from 12800 to 64.

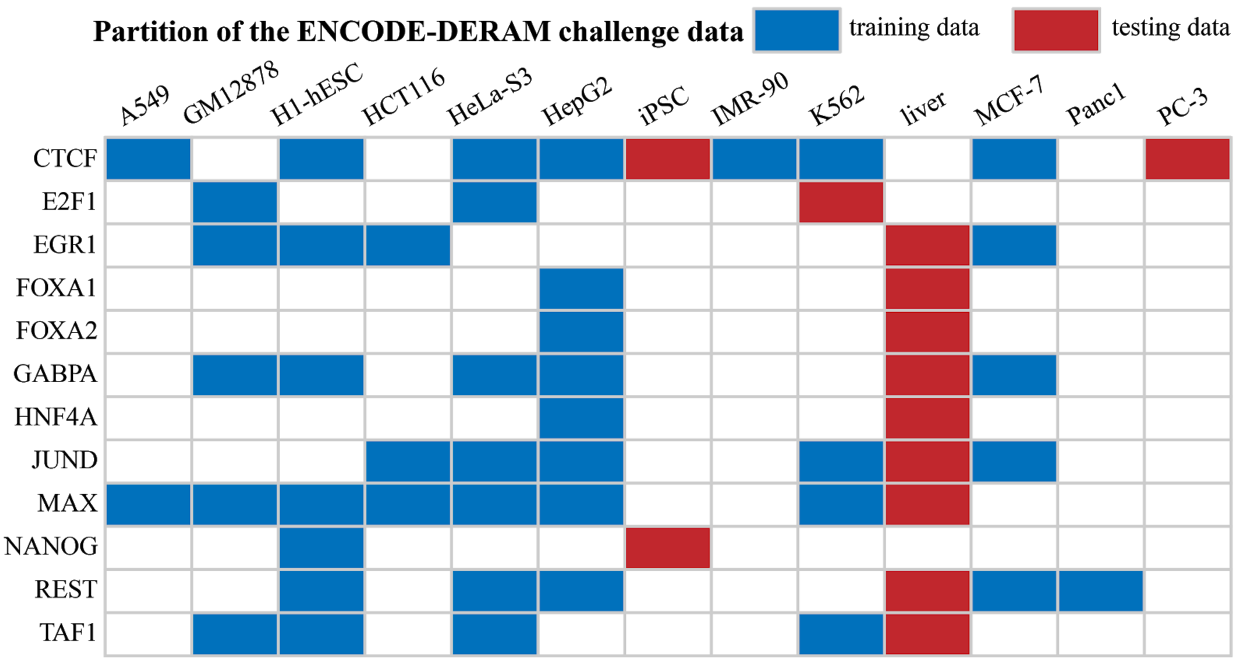


Supplementary Fig. 5: The partition of the dataset used in ENCODE-DREAM *in vivo* Transcription Factor Binding Site Prediction Challenge.

A total of 56 ChIP-seq experiments was used in the ENCODE-DREAM challenge, covering 12 TFs (CTCF, E2F1, EGR1, FOXA1, FOXA2, GABPA, HNF4A, JUND, MAX, NANOG, REST, and TAF1) in 13 cell types (A549, GM12878, H1-hESC, HCT116, HeLa-S3, HepG2, iPSC, IMR-90, K562, liver, MCF-7, Panc1, PC-3). A subset of 43 ChIP-seq experiments were used for model training (blue blocks) and the remaining 13 were held-out for model testing (red blocks)

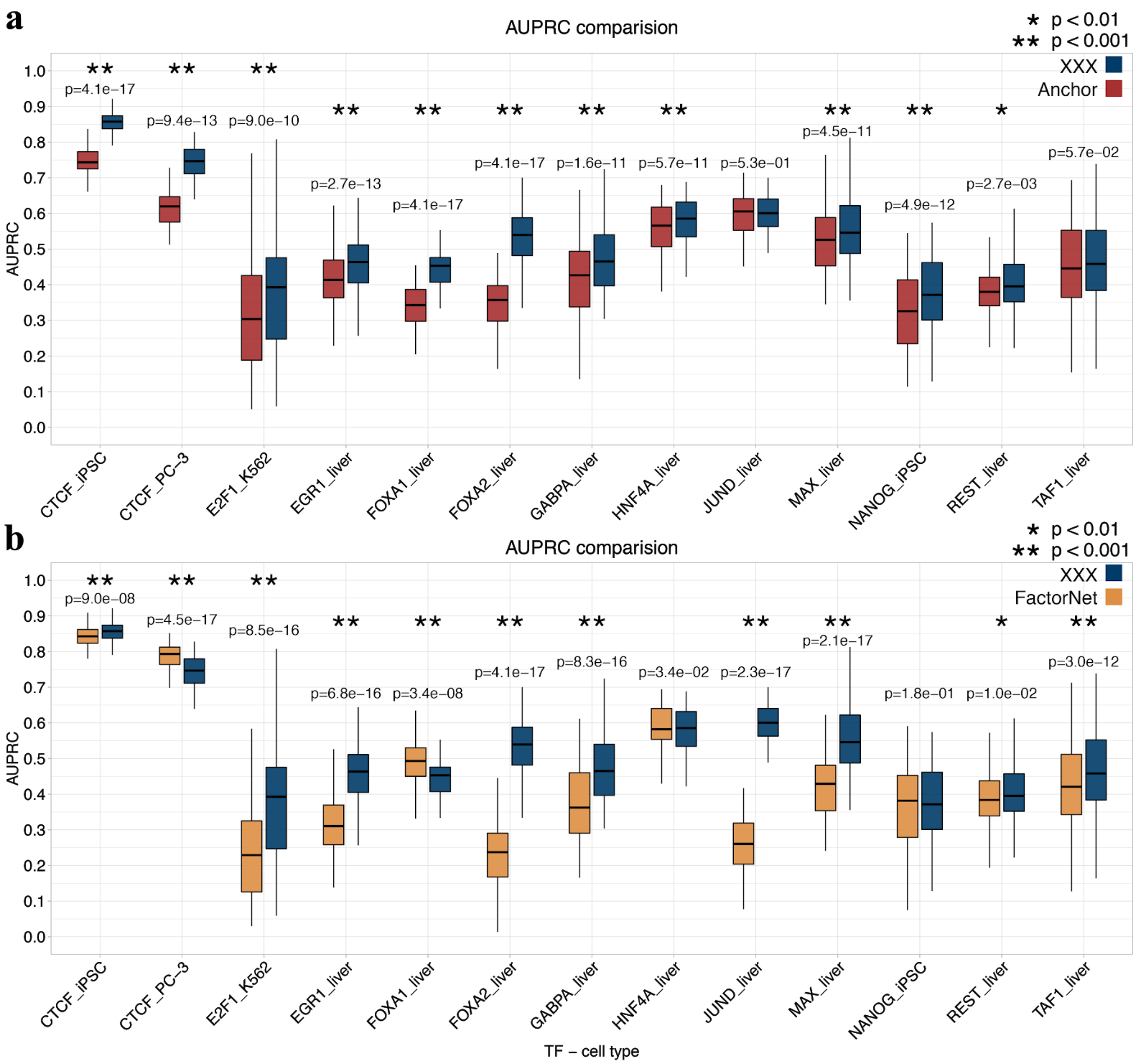


Supplementary Fig. 6: The statistical comparison of Leopard with the state of the art.

For each testing TF-cell type pair, we randomly sampled 100 segments with length of 100kbp and calculated 100 prediction AUPRCs. The paired Wilcoxson signed rank test was performed. Leopard significantly outperformed **a,** Anchor and **b,** FactorNet in the majority of testing TF-cell type pairs. The significant differences are labeled with a single asterisk (p-value < 0.01) or double asterisks (p-value < 0.001).

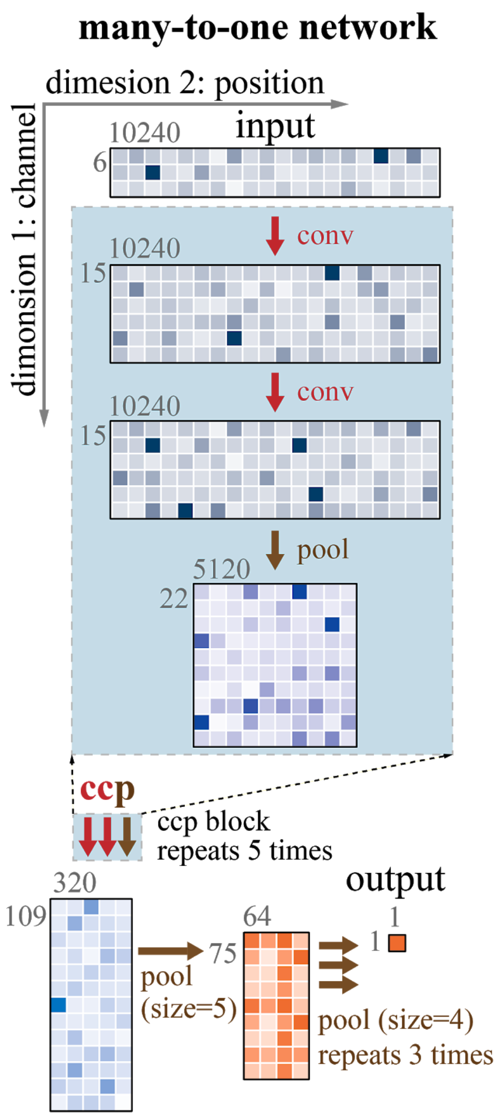


Supplementary Fig. 7: The architecture of the many-to-one model represents a type of convolutional neural network structure commonly used in previous methods.

Compared with Leopard, the many-to-one model only has the encoder (blue) component without the decoder (yellow), representing the common type of convolutional neural network model used in previous methods. At the end of the encoder, multiple max-pooling layers (brown arrows on the right) were used to gradually reduce the size of the output. This type of model is therefore called "many-to-one", since the input size is 10240 (many) and the final output is only 1 value (one).

Supplementary Tables

Supplementary Table 1. The prediction AUROCs and AUPRCs of Leopard in 27 held-out testing TF-cell type pairs.

| TF-cell type pair | AUROC | AUPRC |
|---|---|---|
| CTCF_GM12878 | 0.999 | 0.637 |
| CTCF_HCT116 | 0.994 | 0.516 |
| CTCF_iPSC | 0.997 | 0.661 |
| CTCF_PC-3 | 0.974 | 0.487 |
| E2F1_K562 | 0.997 | 0.256 |
| EGR1_K562 | 0.995 | 0.385 |
| EGR1_liver | 0.995 | 0.265 |
| FOXA1_MCF-7 | 0.959 | 0.358 |
| FOXA1_liver | 0.984 | 0.333 |
| FOXA2_liver | 0.994 | 0.330 |
| GABPA_K562 | 0.998 | 0.429 |
| GABPA_liver | 0.993 | 0.352 |
| HNF4A_liver | 0.962 | 0.430 |
| JUND_A549 | 0.999 | 0.467 |
| JUND_GM12878 | 0.999 | 0.279 |
| JUND_H1-hESC | 0.998 | 0.263 |
| JUND_liver | 0.996 | 0.441 |
| MAX_liver | 0.998 | 0.376 |
| NANOG_iPSC | 0.983 | 0.188 |
| REST_A549 | 0.972 | 0.215 |
| REST_GM12878 | 0.990 | 0.392 |
| REST_K562 | 0.997 | 0.282 |
| REST_liver | 0.979 | 0.214 |
| TAF1_A549 | 0.998 | 0.309 |
| TAF1_HepG2 | 0.994 | 0.409 |
| TAF1_liver | 0.994 | 0.238 |
| **mean** | **0.990** | **0.366** |
| **median** | **0.994** | **0.355** |

Supplementary Table 2. The prediction AUROCs and AUPRCs of Leopard on 23 chromosomes and the overall scores on whole genome when evaluated on the ChIP-exo CTCF binding peaks in HeLa-S3 cell line.

| ChIP-exo | AUROC | AUPRC |
|---:|---:|---:|
| chr1 | 0.989 | 0.309 |
| chr2 | 0.991 | 0.293 |
| chr3 | 0.990 | 0.297 |
| chr4 | 0.993 | 0.294 |
| chr5 | 0.988 | 0.297 |
| chr6 | 0.990 | 0.299 |
| chr7 | 0.989 | 0.296 |
| chr8 | 0.991 | 0.297 |
| chr9 | 0.990 | 0.288 |
| chr10 | 0.992 | 0.304 |
| chr11 | 0.988 | 0.302 |
| chr12 | 0.990 | 0.312 |
| chr13 | 0.992 | 0.284 |
| chr14 | 0.992 | 0.300 |
| chr15 | 0.985 | 0.264 |
| chr16 | 0.992 | 0.285 |
| chr17 | 0.990 | 0.297 |
| chr18 | 0.989 | 0.336 |
| chr19 | 0.990 | 0.300 |
| chr20 | 0.988 | 0.318 |
| chr21 | 0.997 | 0.294 |
| chr22 | 0.996 | 0.301 |
| chrX | 0.994 | 0.292 |
| **overall** | **0.990** | **0.297** |

Supplementary Table 3. The prediction AUPRCs at 200bp resolution of Leopard, Anchor, and FactorNet in 13 held-out testing TF-cell type pairs using the same training and testing data as the ENCODE-DREAM challenge

| TF-cell type pair | Leopard | Anchor | FactorNet |
|---|---|---|---|
| CTCF_iPSC | 0.852 | 0.732 | 0.841 |
| CTCF_PC-3 | 0.716 | 0.593 | 0.774 |
| E2F1_K562 | 0.366 | 0.313 | 0.209 |
| EGR1_liver | 0.440 | 0.400 | 0.291 |
| FOXA1_liver | 0.437 | 0.326 | 0.463 |
| FOXA2_liver | 0.531 | 0.343 | 0.214 |
| GABPA_liver | 0.491 | 0.451 | 0.392 |
| HNF4A_liver | 0.600 | 0.577 | 0.598 |
| JUND_liver | 0.591 | 0.583 | 0.263 |
| MAX_liver | 0.521 | 0.504 | 0.392 |
| NANOG_iPSC | 0.350 | 0.292 | 0.348 |
| REST_liver | 0.414 | 0.380 | 0.385 |
| TAF1_liver | 0.414 | 0.413 | 0.379 |
| **mean** | **0.517** | **0.454** | **0.427** |
| **median** | **0.491** | **0.413** | **0.385** |

Supplementary Table 4. The accession numbers of ChIP-seq data downloaded from the ENCODE project.

| TF-cell type pair | Accession number |
|---|---|
| CTCF_GM12878 | ENCSR000DKV |
| CTCF_HCT116 | ENCSR000BSE |
| EGR1_K562 | ENCSR211LTF |
| FOXA1_MCF-7 | ENCSR126YEB |
| GABPA_K562 | ENCSR000BLO |
| JUND_A549 | ENCSR000BRF |
| JUND_GM12878 | ENCSR000DYS |
| JUND_H1-hESC | ENCSR000BKP |
| REST_A549 | ENCSR892DRK |
| REST_GM12878 | ENCSR000BGF |
| REST_K562 | ENCSR000BMW |
| TAF1_A549 | ENCSR000BPF |
| TAF1_HepG2 | ENCSR000BJN |
